# Contralesional grey matter volume as an index of macrostructural plasticity in patients with brain tumors

**DOI:** 10.1101/2025.06.24.661278

**Authors:** Lucía Manso-Ortega, Sandra Gisbert-Muñoz, Lucia Amoruso, Garazi Bermudez, Santiago Gil Robles, Iñigo Pomposo, Manuel Carreiras, Ileana Quiñones

## Abstract

This research challenges the traditional localizationist view that brain tumors affect only regions directly associated with the lesion, by examining whether they also induce macrostructural alterations in the contralesional hemisphere. We applied Voxel-Based Morphometry, linear regression, and Principal Component Analysis (PCA) to a cohort of 107 adults, including patients with gliomas in the language-dominant left hemisphere and healthy participants. Unlike previous studies, a subset of the clinical population was followed longitudinally for up to four months after oncological treatment, allowing us to describe the temporal progression of structural grey matter changes. Interestingly, a principal component model based on anomaly detection enabled robust differentiation between patients and controls. Patients exhibited significantly greater grey matter volume in the contralesional hemisphere compared to healthy participants, and these structural differences evolved over time, improving the model’s AUC-ROC metrics. Although exploratory, a correlation analysis revealed that these structural changes were negatively associated with postsurgical cognitive performance. Together with the PCA findings, these results suggest that brain tumors induce extensive and dynamic adaptive mechanisms in the contralateral, unaffected hemisphere, likely reflecting altered patterns of structural covariance rather than simple regional volume increases. Understanding whether these changes could represent potential predictors of postoperative cognitive recovery is crucial for developing comprehensive clinical strategies.

**Key Points:**

- Left-hemisphere tumors induce contralesional grey matter increases pre- and post-sugery
- PCA detects altered structural covariance and distinguishes patients from healthy participants
- Contralesional grey matter volume changes correlate with postoperative cognitive performance

**Importance of the Study:** This study challenges the traditional view that brain tumors cause only localized effects by demonstrating widespread macrostructural alterations in the contralesional hemisphere. We reveal increased grey matter volume and altered patterns of structural covariance outside the tumor region. Longitudinal follow-up after surgery shows these changes are dynamic and evolve over time. Further, we identify moderate associations between contralesional grey matter alterations and cognitive performance, suggesting a link between large-scale neuroplastic responses and functional outcomes. These findings offer new insights into tumor-related neuroplasticity and position structural covariance as a promising marker for tracking brain-wide adaptation in this population. The study has translational relevance for developing predictive tools to monitor recovery and guide personalized rehabilitation.

## Introduction

For decades, neuroscience has predominantly focused on studying the brain through a localizationist perspective, examining individual brain regions and their association with cognitive processes in isolation ^1^. However, the emergence of advanced imaging methods has led to a paradigm shift towards recognizing the brain as a complex system organized by distributed and interconnected networks ^2,3^. This evolution emphasizes the need for a holistic perspective to study brain pathology, transitioning to a broad analysis that considers the entire network’s dynamics and their influence on overall brain functionality. This interconnected nature is also one of the reasons why the brain holds great plastic potential and can adapt to the appearance of a tumor, enabling the resection of damaged regions ^4,5^.

In the context of brain tumors, the concept of “metaplasticity” has emerged as a critical area of interest. Metaplasticity refers to the brain’s ability to adjust the threshold of synaptic plasticity based on previous activity, effectively regulating its capacity to undergo plastic changes ^6^. This concept underscores the brain’s dynamic adaptability and suggests that structural changes observed in patients with brain tumors may be part of a broader, network-based response rather than isolated events. While this concept of an interconnected brain has been applied to the study of functional plasticity in patients with brain tumors, demonstrating global effects ^4,5^, structural mechanisms underlying these changes have not been as thoroughly investigated.

Grey matter, which holds a large number of neurons, is essential for information processing. It is also thought to present a higher degree of plasticity than white matter, and its damage is commonly linked to cognitive impairments ^7^. Thus, making grey matter volume (GMV) a good proxy for understanding the macrostructural impact of brain tumors. Methods like Surface-based morphometry (SBM) and Voxel-based morphometry (VBM) facilitate the detection of GMV alterations and contribute to our understanding of morphological changes associated with various neurological conditions. Despite controversies surrounding their application, both SBM and VBM have proven their validity in offering valuable insights to researchers and clinicians alike regarding the morphological landscape of the brain (for a full review of these methods, see ^8^). While SBM measures the brain’s surface characteristics, including cortical volume, thickness, surface area, and gyrification patterns, VBM is the preferred method to focus solely on cortical GMV alterations.

Studies using VBM to explore adaptive changes in contralesional grey matter regions show mixed findings ^9–11^. Some studies report volume increases in these regions, while others observe the opposite pattern. An example can be found in ^9^, they identified a volumetric increase of the contralateral insula compared to that of healthy participants. Similarly, larger volumes were found for the homotopic orbitofrontal cortex ^12^, temporal lobe ^13^, frontal lobe ^14^ and the cerebellum ^15^. In contrast, significantly smaller volumes in the medial temporal cortex contralateral to the lesion has also been observed ^11^. These discrepancies have been associated with differences in methodological approaches and are influenced by factors such as tumor location, type, and patient characteristics ^10^.

Previous evidence from studies using either VBM or SBM show that structural changes within certain networks are consistent regardless of tumor location ^5,16,17^. For instance, using SBM, another study found volume increases in both hemispheres, including regions such as the right cuneus, left thalamus, and right globus pallidus at a superficial level, which correlated with lesion size ^14^(see also ^5^). Decreased cortical thickness and fractal dimension were also found regardless of tumor volume, location or grade ^10^, reflecting the interconnected nature of brain networks ^18^.

Despite the valuable contributions made by these studies, there is a notable gap in the literature regarding the longitudinal progression of GMV changes in patients with brain tumors. To date, only two studies show GMV changes over time for patients with brain tumors. In the study by ^19^, in comparison to healthy participants, patients with left-hemisphere tumors showed larger GMV in the right inferior temporal gyrus (ITG) and right temporal pole (STG), while patients with right hemisphere-tumors showed more volume in left ITG. Post-operatively, in relation to healthy participants, patients with right tumors presented increased GMV in the left middle temporal gyrus (MTG), which was positively correlated with memory and visuospatial abilities ^19^. No differences were found by comparing patients before and after surgery. The other study exploring the temporal progression of said changes is restricted to a pediatric population ^20^, limiting the generalization of the results. They combined SBM and VBM and reported increased GMV in the contralateral hemisphere six months post-surgery, particularly in the frontal lobe. This increase was correlated with enhanced functional activity and interpreted as a compensatory mechanism to support high-order cognitive functions. Given that neuroplasticity is likely to be constrained by developmental factors, further research is essential to determine how these findings translate to the adult brain.

Building from this background, the present study aims to advance the understanding of the complexity of structural plasticity mechanisms by examining the entire contralesional hemisphere in patients with brain tumors through VBM. Specifically, we addressed three main objectives. First, we assessed macrostructural differences in the contralesional hemisphere between patients with left-hemisphere tumors and healthy participants. We hypothesized widespread macrostructural alterations would differentiate patients from healthy participants, reflecting the brain’s adaptive response to the tumor presence. We further anticipated that interconnected brain regions would exhibit similar GMV changes, illustrating the impact of a brain tumor on remote areas and providing evidence for the brain’s networked organization. Second, we tracked the progression of structural changes over time by comparing pre-surgical and three-month post-surgical imaging data. Third, we examined the relationship between volumetric changes in the contralesional hemisphere and cognitive performance. We predicted that alterations in GMV of the contralesional hemisphere would be associated with preserved cognitive abilities, which could reflect a compensatory mechanism. The brain’s functional network organization might allow for neuroplastic compensation, facilitating cognitive resilience and enabling safe resection of tumor-affected areas. A comprehensive understanding of structure-behavior coupling processes could inform treatment strategies aimed at mitigating cognitive impairments.

## Methods

### 1. Participants

#### 1.1. Patients

A total of 37 patients (24 males, mean age = 45.16 years, SD = 13.81) diagnosed with brain tumors within the left hemisphere participated in this study. Patients were recruited via the Hospital Universitario Cruces Bilbao (Spain), where they received their diagnosis and underwent awake brain surgery for tumor resection (M.D. Garazi Bermudez, MD. Ph.D. Santiago Gil-Robles, and M.D. Iñigo Pomposo Gastelu—as the neurosurgeons in charge). Initial neurological evaluation showed no motor, somatosensory, or linguistic deficit. All patients had normal hearing and normal or corrected-to-normal vision. In Table 1, we present a summary of the tumor characteristics, such as tumor grade and tumor type according to affected cell type (Johnson et al., 2012). Further, we include the post-surgical assessment of 21 of those patients, 3 months after surgery, including neuropsychological and demographic variables.

**Table 1.** Description of tumor characteristics (n = 37) based on the 2021 World Health Organization (WHO) ^21^ pathophysiological classification.

| **Overall tumor classification** |  |
| --- | --- |
| High-grade glioma | 15 (41%) |
| Low-grade glioma | 19 (51%) |
| Other | 3 (8.1%) |
| **Tumor classification (affected cell type)** |  |
| Glioblastoma | 6 (16%) |
| Astrocytoma | 16 (43%) |
| Oligodendroglioma | 10 (27%) |
| Oligoastrocytoma | 1 (2.7%) |
| Pleomorphic Xanthoastrocytoma | 1 (2.7%) |
| Dysembryoplastic Neuroectodermal Tumor (DNET) | 2 (5.4%) |
| Cortical Dysplasia* | 1 (2.7%) |

All included tumors induce neuroplastic changes due to their impact on neural tissues. Extra-axial lesions were excluded from this study.
\*Not considered in the WHO classification as it is not a neoplastic lesion.

#### 1.2. Healthy participants

As a healthy control group, we incorporated a cohort of 70 individuals matched in terms of age and sex (42 women, 28 males, mean age = 45.23 years, SD = 12.41). We included them to characterize the structural relationships within the typical healthy brain, enabling us to discern any potential variations among the patient cohort. This data was included as a control group in a previous study about structural organization in patients with brain tumors, see ^16^.

### 2. Procedure

Patients completed an initial screening approximately one week before surgery, in which MRI, fMRI, and behavioral data were collected. After surgery, patients completed a post-surgical screening after approximately 3 months (mean days between surgery and first post-surgical screening = 128.63, SD = 47.62); the exact count of days between the surgery and the post-surgical screening for each patient can be found in the Supplementary Material. Healthy participants completed a single time-point data collection including MRI, fMRI, and behavioral data.

At the beginning of each session, participants received instructions about the techniques and tasks to perform. Informed consent was acquired from all participants prior to data collection. Data was collected at the Basque Center on Cognition, Brain and Language (BCBL). Healthy participants were economically compensated for their time. Patients’ and healthy participants’ data collection was approved by the Ethics Board of the Euskadi Committee and the Ethics and Scientific Committee of the Basque Center on Cognition, Brain, and Language, BCBL (protocol code PI2020022, date of approval: 26 May 2020 and protocol code: 270220SM, respectively). The study protocol was conducted in accordance with the Declaration of Helsinki for experiments involving humans.

#### 2.1. Cognitive Assessment

All participants completed a set of standardized neuropsychological and behavioral evaluations. For patients, the cognitive assessment was completed approximately one week before surgery and 3 months after the intervention, allowing for a comparison of cognitive performance before and after surgery. For healthy participants, this evaluation was only conducted once.

Given the functional relationship between the left hemisphere—where all patients in the clinical sample exhibited damage—and language, the cognitive assessment primarily focused on three domains: language, working memory, and intelligence (verbal and non-verbal) (see Table 2). To obtain a measure for each domain, we constructed composite variables by aggregating the standardized performance values across related tasks. Composite variables combine multiple variables ^22^, proving valuable in clinical contexts for handling missing data and capturing the multidimensional nature of cognitive measures ^23^. To enable the combination of the cognitive tasks, we computed z-scores for all participants. This transformation standardizes the data, representing each data point’s deviation from the mean of the healthy participants. The z-score calculation followed the formula: *z-score = (x-μ)/σ,* where *μ* is the mean and *σ* the standard deviation (SD). We used the mean and SD of the healthy participants as the reference for patients. Missing values were excluded on a task-by-task basis when computing composite variables, meaning that participants were included in a composite score as long as they had data for at least one of the task in that domain (see Supplementary Material Table 1 and Table 2 for full details on missing data patterns).

**Table 2.**
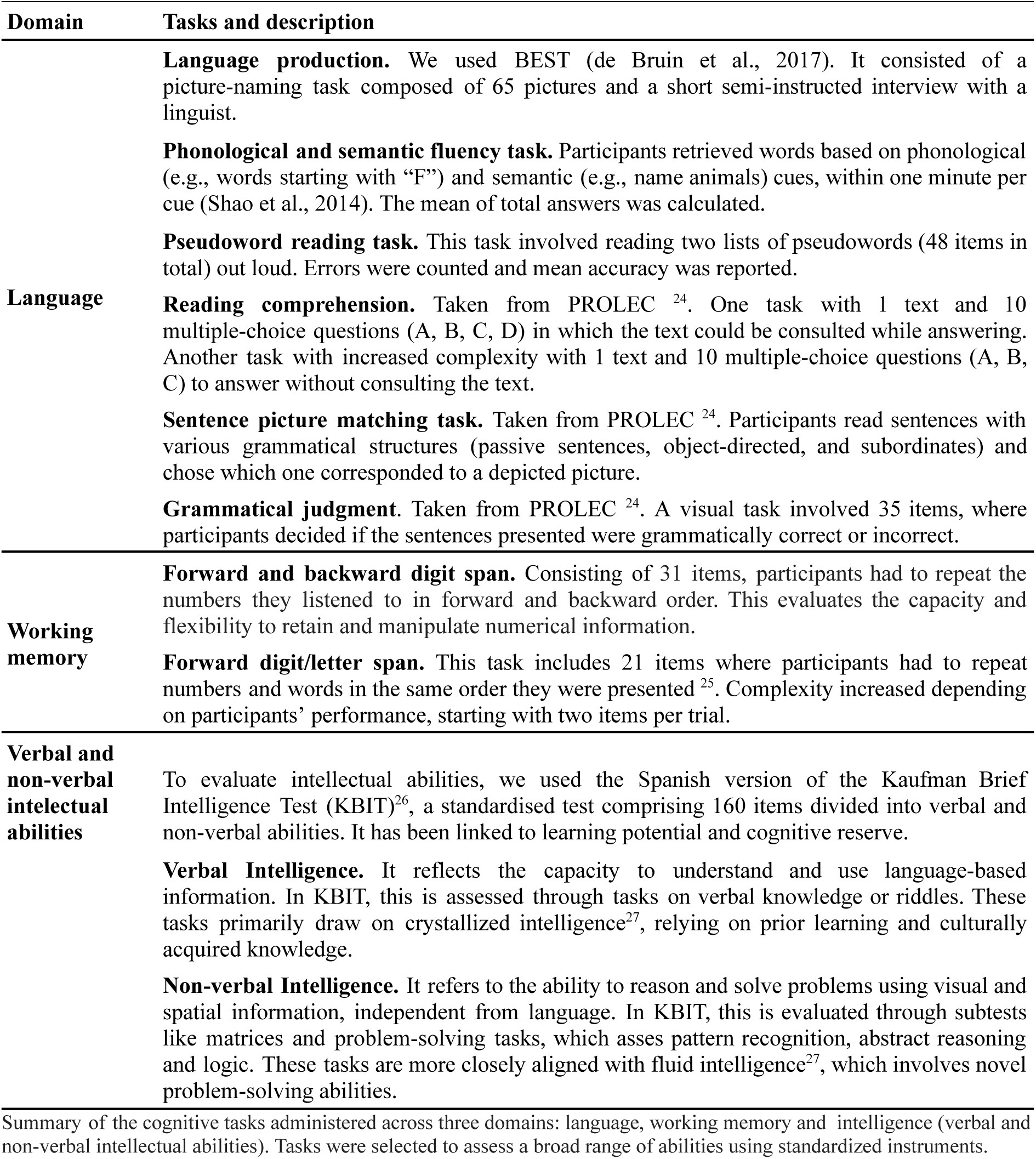
Cognitive assessment.

#### 2.3. MRI Data Acquisition

Whole-brain MRI data were acquired using a 3T Siemens Magnetom Prisma Fit scanner (Siemens AG, Erlangen, Germany). To obtain high-resolution images, a 3D ultrafast gradient echo (MPRAGE) pulse sequence was employed, using a 64-channel head coil. The T1-weighted images were acquired with the following parameters: 176 contiguous sagittal slices, voxel resolution of 1×1×1 mm³, repetition time (TR) = 2530 ms, echo time (TE) = 2.36 ms, 256 image columns, 256 image rows and a Flip angle (Flip) = 7°. Similarly, the T2-weighted images were acquired with 176 contiguous sagittal slices, voxel resolution of 1×1×1 mm^3^, TR = 3390 ms, TE = 389 ms, 204 Image columns, 256 Image rows and Flip = 120°. For each participant, the origin of T1 and T2 weighted images for each pre and post-surgery session were adjusted to the anterior commissure. Subsequently, all structural images were aligned using co-registration techniques.

#### 2.4. Lesion reconstruction for patients with brain tumors

The lesion-affected area for each patient was reconstructed by two experts under the supervision of the senior author (I.Q.) and the neurosurgeon in charge of the patient’s awake craniotomy (G.B.). These masks were manually drawn slice by slice on the native space using the free and open-source software MRIco-GL ^28^. To accurately determine the extent of the lesion, we employed a combination of the co-registered T1- and T2-weighted images and applied multiple intensity thresholds. The lesion was then normalized to the Montreal Neurological Institute (MNI) template and the alignment between the reconstructed lesion and the lesion in the native space was checked. For each patient at each time point, a volume of interest (VOI) was created and smoothed. This VOI was used to estimate the tumor volume (cm3) and the extent of resection. The extent of resection (cm3) was estimated on postoperative imaging as (Volume of preoperative 3D Tumor Reconstruction / postoperative resection) * 100/preoperative tumor volume)). A 3D reconstruction of the overlap lesion map by location is depicted in Figure 1.

**Figure 1.**
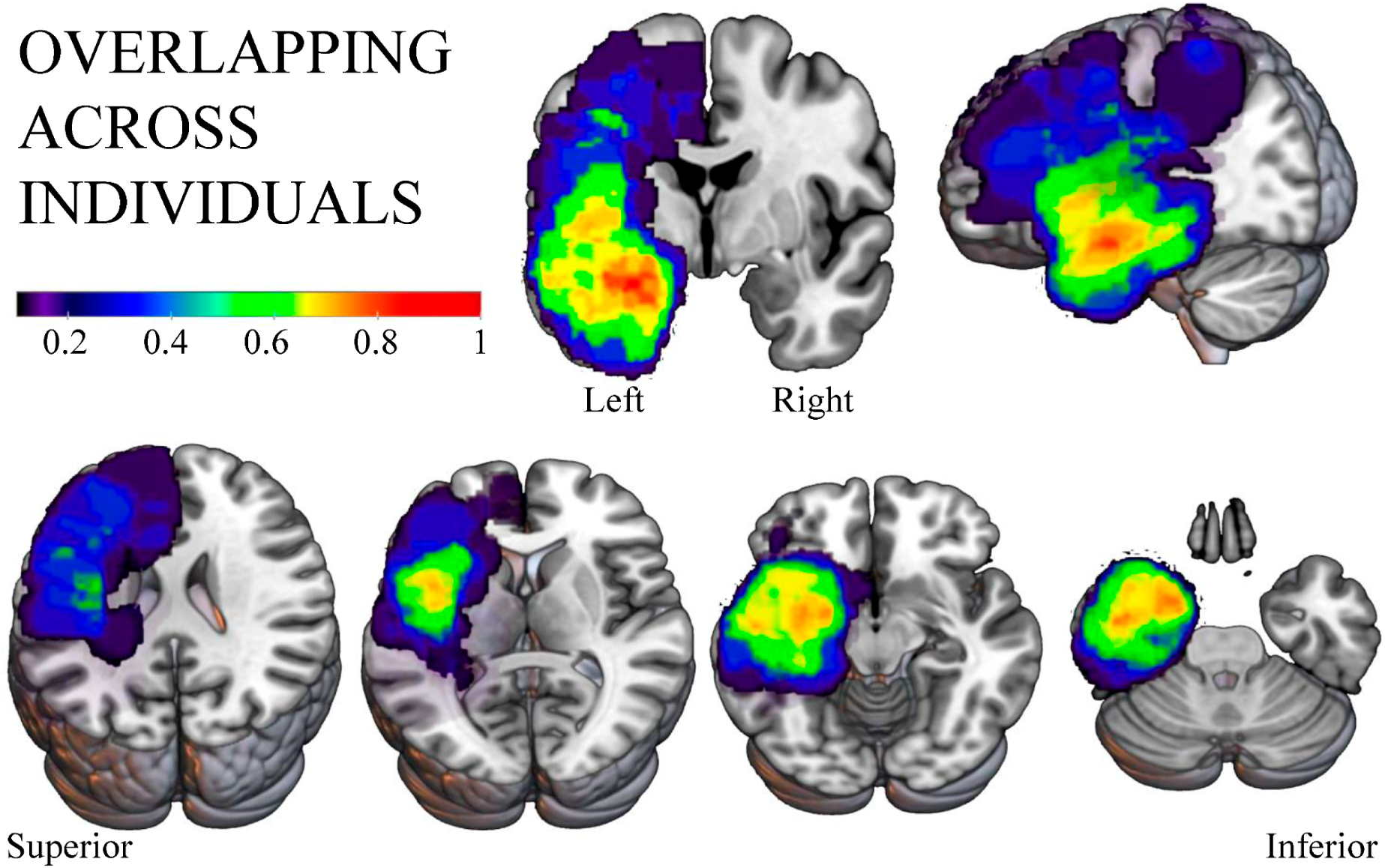
Lesion overlap per tumor location (n = 37). The heatmaps—from red to blue—represent the number of overlapping tumors within groups in percentages. We represent 16 frontal, 16 temporal, and 5 parietal lesions.

### 2.5. MRI Analysis

#### Voxel-based morphometry analysis (VBM)

To measure GMV for healthy controls and patients at each time point, we performed a VBM analysis using SPM12 in MATLAB 2014b. VBM is a semi-automated neuroimaging method used to quantify macrostructural changes in the brain, such as changes in volume or density. Prior studies have demonstrated its efficacy in various contexts, from development to pathological populations ^29^, facilitating the understanding of structural brain alterations, and inferring the presence of tissue atrophy or, contrarily, tissue expansion. In brief, VBM conducts statistical tests across all voxels within images and segments them into different tissues to enable identifying differences in brain anatomy between groups or across time.

For the current study, the preprocessing pipeline was standardized for both groups and contained specific steps to account for the lesion in the case of patients. Furthermore, the pipeline used for the current VBM analysis takes into account any possible distortion that could be caused by the normalization algorithm in pathological brains ^30^.

For all participants, first, we performed a manual trimming of the T1 and T2 weighted images to remove non-relevant background and visually inspected them to check for artifacts or distortions in image acquisition. Then, images were manually re-oriented and shifted to set the origin at the anterior commissure, a bundle of white matter fibers that connect the anterior lobes of the brain. To match images spatially, we coregistered them setting the T1 as the reference to improve the segmentation. In the case of post-surgical time points for patients, we also coregistered these images to the pre-surgical time point T1 as a reference. Then, using the information from the T1 and T2 images, T1 MPRAGE-weighted images were segmented into grey matter (GM), white matter (WM), and cerebrospinal fluid (CSF) through the segmentation module in SPM12. Segmented images were smoothed with an 8 x 8 x 8 Gaussian kernel to improve statistical inferences. Before estimating the volumes, for patients, we manually created a lesion mask in MicroGL ^28^. For the post-surgical time points, the tumor mask included both the resection and tumor remains if any. We binarised this mask, normalized it to the MNI space, resliced it to the T1 space, and then resliced it to the normalized modulated space. Then, for the patients, we subtracted the tumor masks to account for the presence of the lesion and excluded it from further analysis. We computed the total intracranial volume (TIV) for each participant using the volumes of the native segmentations (GM, WM, and CSF).

As a spatial constraint for all participants, we used the automated anatomical labeling atlas (AAL2) ^31,32^ to obtain GM volume per region. This atlas comprises 116 regions of interest (ROIs). Through in-house Matlab codes, we converted all ROIs from the atlas into the native space of each participant using the transformation matrix (iy) obtained during segmentation to avoid further transformations to the patient’s images. We binarised the ROIs to create masks and then resliced them to the T1 space. We estimated the volume of each ROI in the native space of the participant and obtained the volume of each ROI (in the case of patients, excluding the tumor mask). Each of the 116 ROIs of the AAL atlas for each participant was divided by the total intracranial volume (TIV), resulting in proportionally scaled scores according to ROI Volume/TIV * 1000 (mm), instead of using the raw GM segmentations. This reduces potential bias due to individual variabilities in brain sizes and enables comparisons across different participants and populations. A detailed diagram of the VBM process can be found in **Figure 2**.

**Figure 2.**
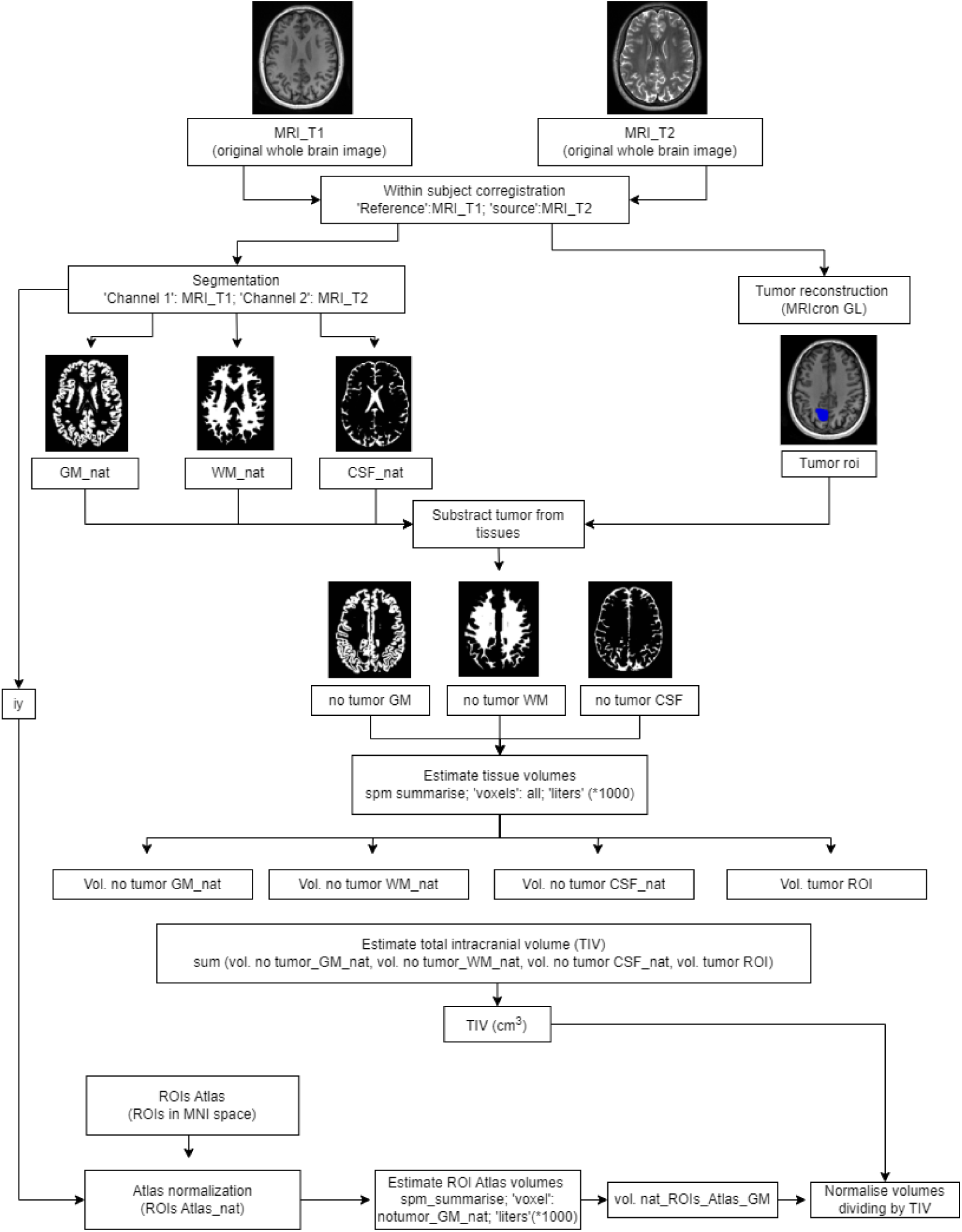
Procedure followed for the VBM analysis of patients’ data. For healthy participants, the same procedure was followed, excluding the steps directly related to handling the tumor mask. For this structural analysis, we developed codes using MATLAB (2014b release, Mathworks, Inc., Natick, MA, USA), functions from Statistical Parametric Mapping (SPM12, Welcome Department of Cognitive Neurology, London, UK), and related toolboxes. Codes are public and available on GitHub (see Data availability statement).

### 2.6. Linear regression analysis

A linear regression analysis was used to evaluate the direction of differences on right hemisphere volume between patients and healthy participants while accounting for the influence of demographic factors such as age and sex. We generated a single metric of right hemisphere volume by aggregating data from 54 ROIs from the AAL2 atlas, which was included as the dependent variable in our analysis. This analysis was performed using the lme4 package in R. This method allowed us to evaluate main effects and interactions, offering insights into how these factors may modulate structural brain changes.

### 2.6. Principal Component Analysis (PCA)

Principal Component Analysis (PCA) is a versatile machine learning method that transforms a dataset into a set of linearly uncorrelated variables called principal components. These principal components represent the directions along which the variance of the data is maximized. The first principal component captures the largest possible variance, while each succeeding component captures the highest variance possible under the constraint that it remains orthogonal to the preceding components. PCA serves multiple purposes: it can simplify a large dataset into a smaller one while preserving essential patterns and trends, extracting key features weighted and ordered by their relevance ^33^. Additionally, PCA can be used for anomaly detection by identifying data that deviate significantly from a pattern learned from normal training data used to calculate the principal components ^34^. This is achieved by measuring the reconstruction error, with data points exhibiting high reconstruction error considered anomalies, as they do not fit well into the structure defined by the training data.

**Objective I).** *Assess whether there are macrostructural differences in GMV between patients and healthy participants that extend within the contralesional hemisphere*. To address this goal, we selected 54 ROIs from the initial 116 extracted. First, we performed a linear regression analysis to examine volumetric differences accounting for the effects of age, sex and group and their interactions in the total volume of the right hemisphere. The measure of volume was the sum of the 54 regions selected. Further, we complemented this analysis with PCA as an anomaly detection approach. We normalized the data using StandardScaler in Sklearn ^35^, scaling each ROI to be zero-centered with a standard deviation of 1, allowing for equal consideration of all ROIs regardless of size. Then, we divided our dataset into training and test. We trained the model using 54 ROIs from the right hemisphere of 40 randomly selected healthy participants from our initial sample of 70. To ensure reproducibility, a seed was incorporated during the random selection process. The trained model encapsulates the structural characteristics of the healthy right hemisphere as a linear combination of features, thereby providing a reference.

For the trained model, we calculated the variance explained by each principal component. For the test phase, we included the remaining 30 healthy participants and our sample of 37 patients. With the reconstruction error, this test aimed to evaluate if the patient’s data deviates from the normal structure of the right hemisphere. Successful differentiation between healthy participants and patients during the test phase indicates significant differences in GMV across groups. We evaluated the model’s performance using the area under the ROC curve (AUC), which summarizes the model’s ability to distinguish between patients and healthy participants based on volumetric data.

**Objective II).** *Assess differences in GMV in patients across time*. We used the same training as in the previous section as a reference for the normal structure of the right hemisphere. For the test phase, we included 30 healthy participants, 37 patients at the pre-surgical time point, and 21 patients at the post-surgical timepoint. This testing phase focused on evaluating whether patients at the post-surgical time point differ more from healthy participants compared to patients at the pre-surgical timepoint, thereby assessing changes in GMV over time. We used the reconstruction error as a measure of deviation from the normal structure and evaluated the model’s performance using the area under the ROC curve (AUC).

**Objective III).** *Understand the relationship between cognitive performance over time and GMV changes*. We evaluated cognitive performance progression (Performance After Surgery - Performance Before Surgery) to obtain a measure of the amount and direction of the change in cognitive performance, where positive values represent an improvement in cognitive performance. We included a measure of progress on language, working memory, and intelligence. We used robust correlations to explore potential associations between cognitive progression and right volumetric variation. Specifically, we used percentage-bend correlations (Wilcox, 1994, 2016), which provide an accurate estimate of the true relationship between two variables while reducing the influence of outliers. To obtain distribution-free significance estimates, *p*-values were derived from 500 random permutations. Analyses were performed with the Pingouin Python package.

## Results

### *I.* Patients with left-hemisphere tumors exhibit macrostructural differences in the contralesional hemisphere compared to healthy participants

The linear regression model explained a substantial proportion of the variance of the volume in the right contralesional hemisphere (R^2^ = 0.651, adjusted R^2^ = 0.651), indicating that 65.1% of the variance in volume values was accounted for by the predictors. The model was statistically significant (F(6, 5771)) = 1795, *p* < 0.001). As expected, age was a strong negative predictor of right hemisphere volume (Estimate = 32.155, *p* < 0.001), indicating that volume decreases with age. There was also a significant effect of sex, with males showing larger left hemisphere volumes compared to females (Estimate = 32.155, *p* < 0.001). Patients exhibited significantly higher right hemisphere volume than healthy participants (Estimate = 42.226, *p* < 0.001).

A significant interaction between age and sex was observed (Estimate = -0.441, p < 0.001), suggesting that the rate of volume decline with age is steeper in males than in females. The interaction between age and group was also significant (Estimate = -0.903, *p* < 0.001), indicating that patients exhibit a more pronounced age-related decline in right hemisphere volume compared to healthy participants. There was a smaller but significant interaction between sex and group (Estimate = -2.722, *p* = 0.0036), suggesting that sex differences in volume may vary slightly between patients and controls.

**Figure 4.2.**
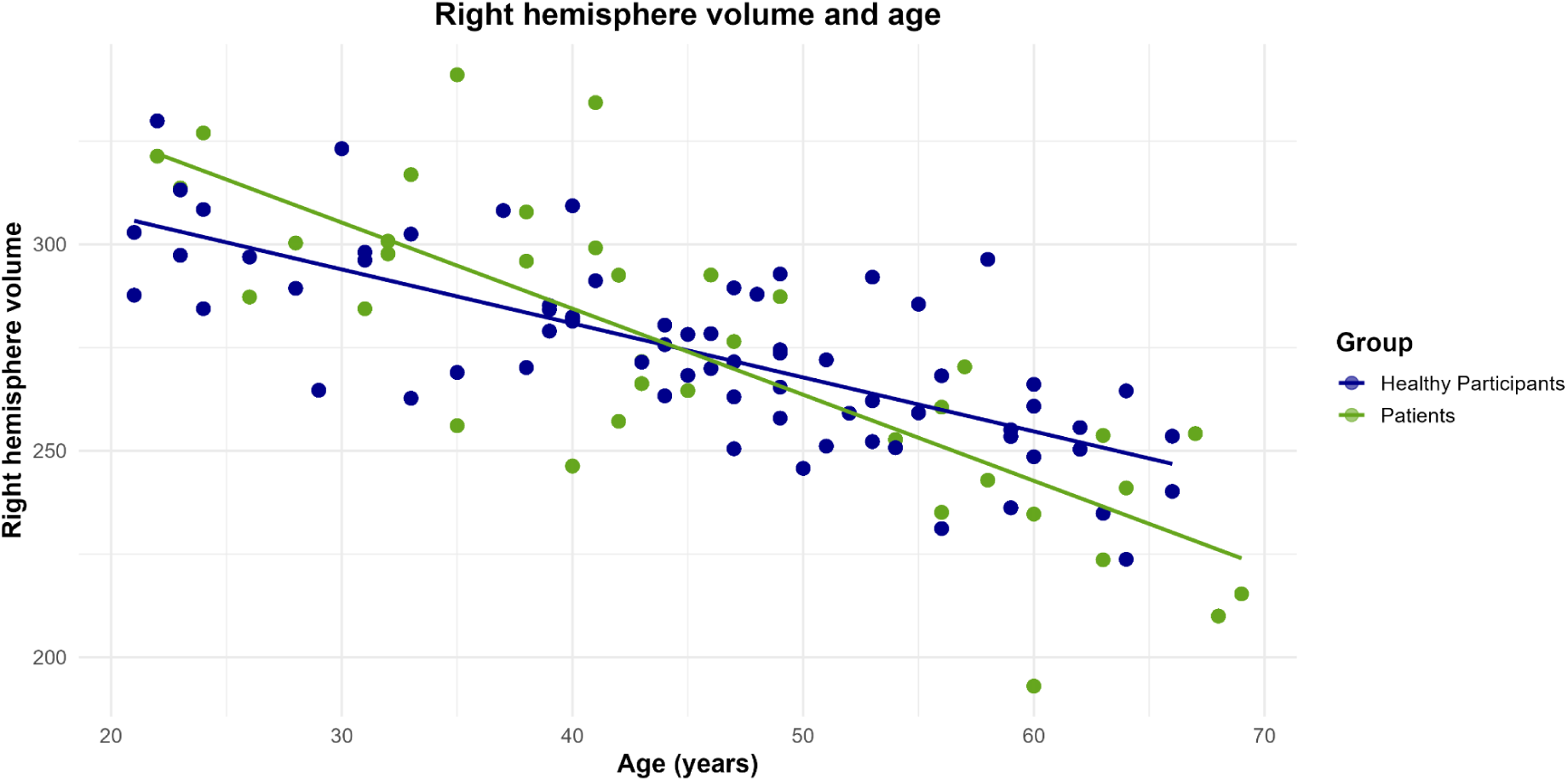
The plot depicts the relationship between age (in years) and right hemisphere volume across two groups: Healthy Participants (dark blue) and Patients with left-hemisphere tumors (green). The x-axis represents participants’ age, and the y-axis shows their right hemisphere volume. Solid trend lines represent linear regressions for each group.

**Table 4.3.**
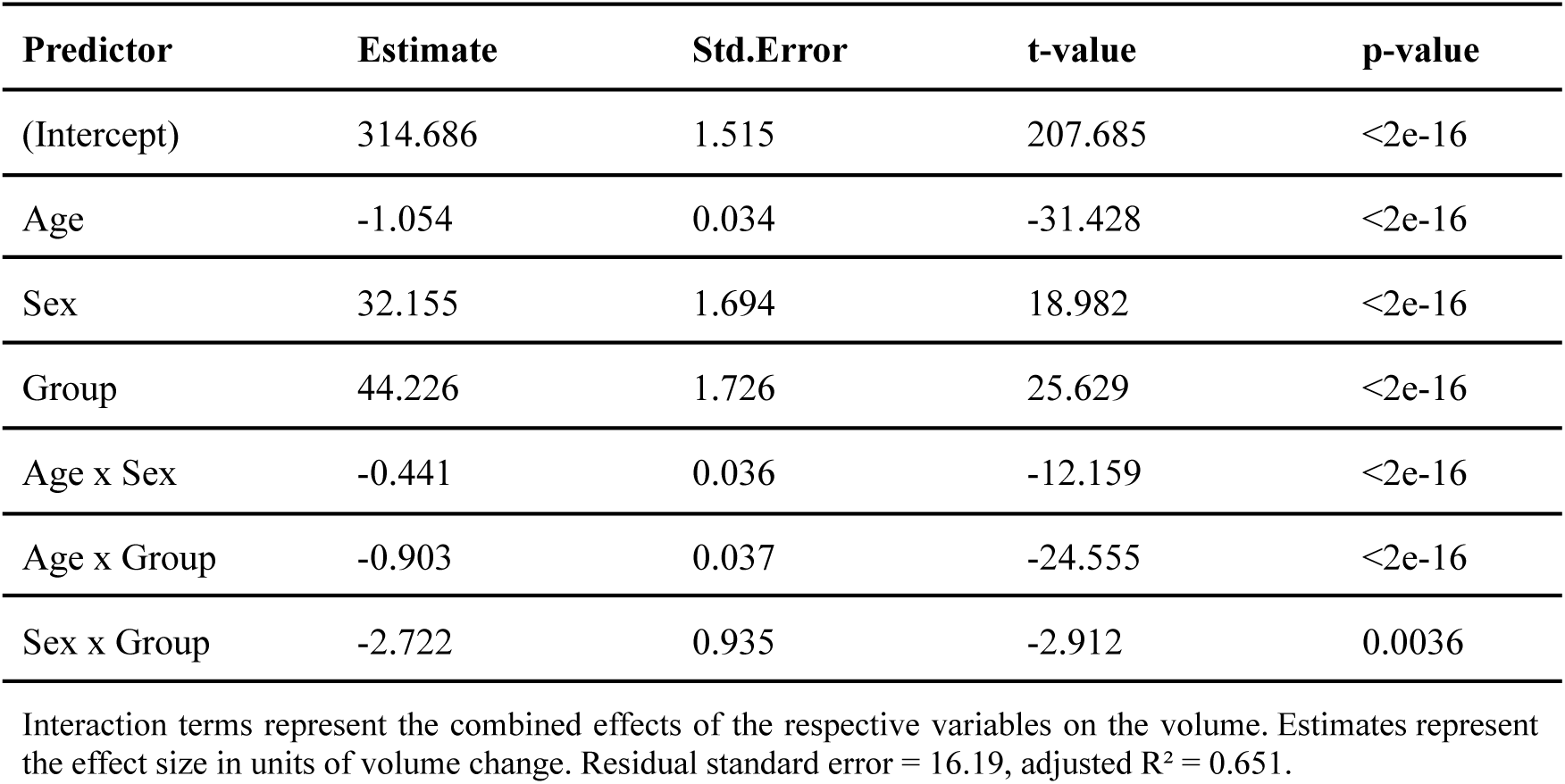
Regression analysis of right hemisphere volume.

| Predictor | Estimate | Std.Error | t-value | p-value |
| --- | --- | --- | --- | --- |
| (Intercept) | 314.686 | 1.515 | 207.685 | <2e-16 |
| Age | -1.054 | 0.034 | -31.428 | <2e-16 |
| Sex | 32.155 | 1.694 | 18.982 | <2e-16 |
| Group | 44.226 | 1.726 | 25.629 | <2e-16 |
| Age x Sex | -0.441 | 0.036 | -12.159 | <2e-16 |
| Age x Group | -0.903 | 0.037 | -24.555 | <2e-16 |
| Sex x Group | -2.722 | 0.935 | -2.912 | 0.0036 |

Interaction terms represent the combined effects of the respective variables on the volume. Estimates represent the effect size in units of volume change. Residual standard error = 16.19, adjusted $R^2 = 0.651$ .

Further, with the PCA model we trained using a sample of healthy participants, we used the eigenvalues to evaluate how many components are needed to explain the training data. For the training dataset, the cumulative variance plot shows we need at least 12 components to explain 90% of the variance (see Figure 3B). However, from the fourth component onwards, each component explains less than 0.05 of the variance. As expected, in Figure 3C, we observe that with only the scores of the first 2 components, we are not able to differentiate between groups. In an anomaly detection approach, we do not need to use all the data but rather select which characteristics are important to represent it. According to the variance explained in Figure 3A and to keep the number of components small and avoid overfitting, we combined the reconstruction error of the first 3 components (0.67 of the variance explained). Within the test phase, we computed the reconstruction error for each participant, reflecting the degree of inaccuracy in distinguishing between healthy participants and patients.

**Figure 3.**
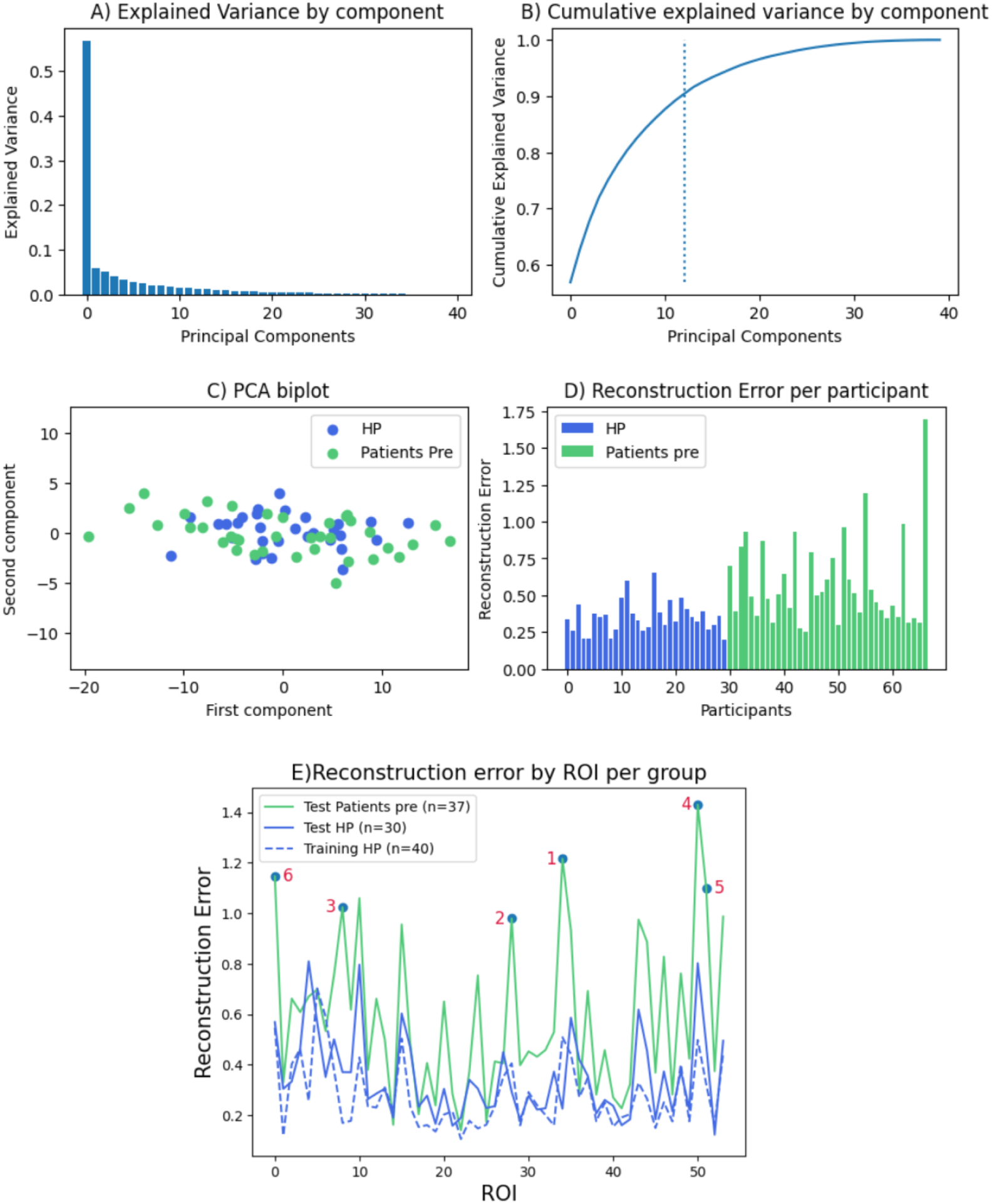
A) Variance explained by each component in the training dataset. The Y-axis represents the variance explained and the X-axis represents the number of the component. B) Cumulative variance explained by the components in the training dataset. The Y-axis represents the variance explained, and the X-axis represents the number of the component. C) Test data represented as the first two components. The first component is the Y-axis and the second component is the X-axis. D) Reconstruction error for each participant in the test phase. The Y-axis represents the reconstruction error values, and the X-axis each of the participants. E) Reconstruction error for each ROI in each of the groups. Y-axis reconstruction error values, X-axis right hemisphere ROIs. Each line represents one of the groups. 1) Olfactory, 2) Hippocampus, 3) Cerebellum 7, 4) Temporal Pole Mid, 5) Temporal Pole Sup, and 6) Amygdala

From this reconstruction error per participant plot (Figure 3D), we can observe that our model performs worse when reconstructing the data for patients compared to healthy participants. To objectively assess the model’s classification performance, we calculated the area under the ROC curve (AUC-ROC). We obtained an AUC-ROC value of 0.78, indicating good discrimination between the groups based solely on the GMV of the right hemisphere. These results suggest that patients and healthy participants do differ in the structure of the overall right hemisphere, with certain ROIs being more different across groups. Additionally, we extracted the reconstruction error for each of the 54 ROIs. We focused on ROIs that were difficult to reconstruct for patients but not for controls to minimize errors arising from the model itself (reconstruction error for the pre-surgical time-point - reconstruction error for healthy participants). The specific ROIs with notable reconstruction errors in patients compared to healthy participants are as follows: 1) Olfactory (0.17), 2) Hippocampus (0.08), 3) Cerebellum 7 (0.08), 4) Temporal Pole Mid (0.07), 5) Temporal Pole Sup (0.07) and 6) Amygdala (0.06), as depicted in Figure 3E. We found that our model had higher reconstruction errors in the majority of ROIs for the patient group compared to healthy participants in both the training and test phases.

### *II*. Macrostructural differences in GMV for patients with left-hemisphere tumors progress over time

To explore grey matter changes over time, specifically 3 months after surgery, we used the same approach and model as in the previous analysis but in the test phase we also included 21 patients with data before and after surgery and we limited the number of patients in the pre-surgical time point to those same 21 patients. We found that the reconstruction error is higher in the post-surgical time point, which suggests that changes in GMV evolve over time.

The ROC-AUC value was 0.93, indicating a very high performance in discriminating between patients’ right hemisphere ROIs after surgery and healthy participants. The ROIs with the highest reconstruction error for patients in the post-surgical timepoint were: 1) Cerebellum 8 (0.41); 2) Cerebellum 7b (0.15); 3) Frontal Med Orb (0.09); 4) Amygdala (0.05); 5) Frontal Sup Orb (0.04); and 6) Temporal Pole Sup (0.03).

### *III.* GMV of the contralesional hemisphere could be a predictor of cognitive performance after surgery

The relationship between GM volume in the contralesional right hemisphere and progress in cognitive performance was tested for three different domains: language (n=15), working memory (n=10), and intelligence (n=20). A representation of the progress in each cognitive variable for each participant can be seen in Figure 4.5.B.

**Figure 4.**
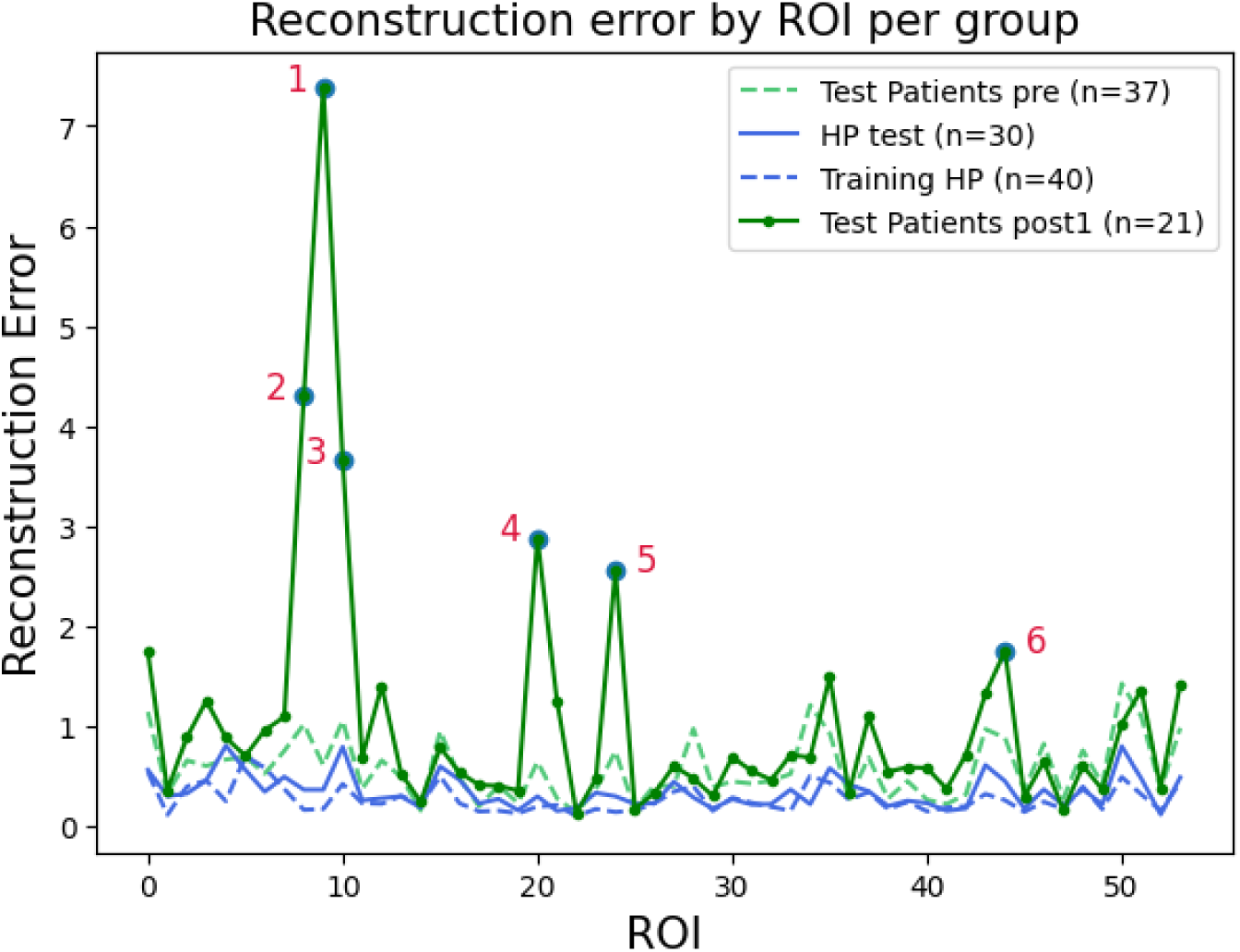
Reconstruction error for each ROI in each of the groups from the test phase in comparison to the training data. Y-axis reconstruction error values, X-axis right hemisphere ROIs. Each line represents one of the groups.

**Figure 4.5.**
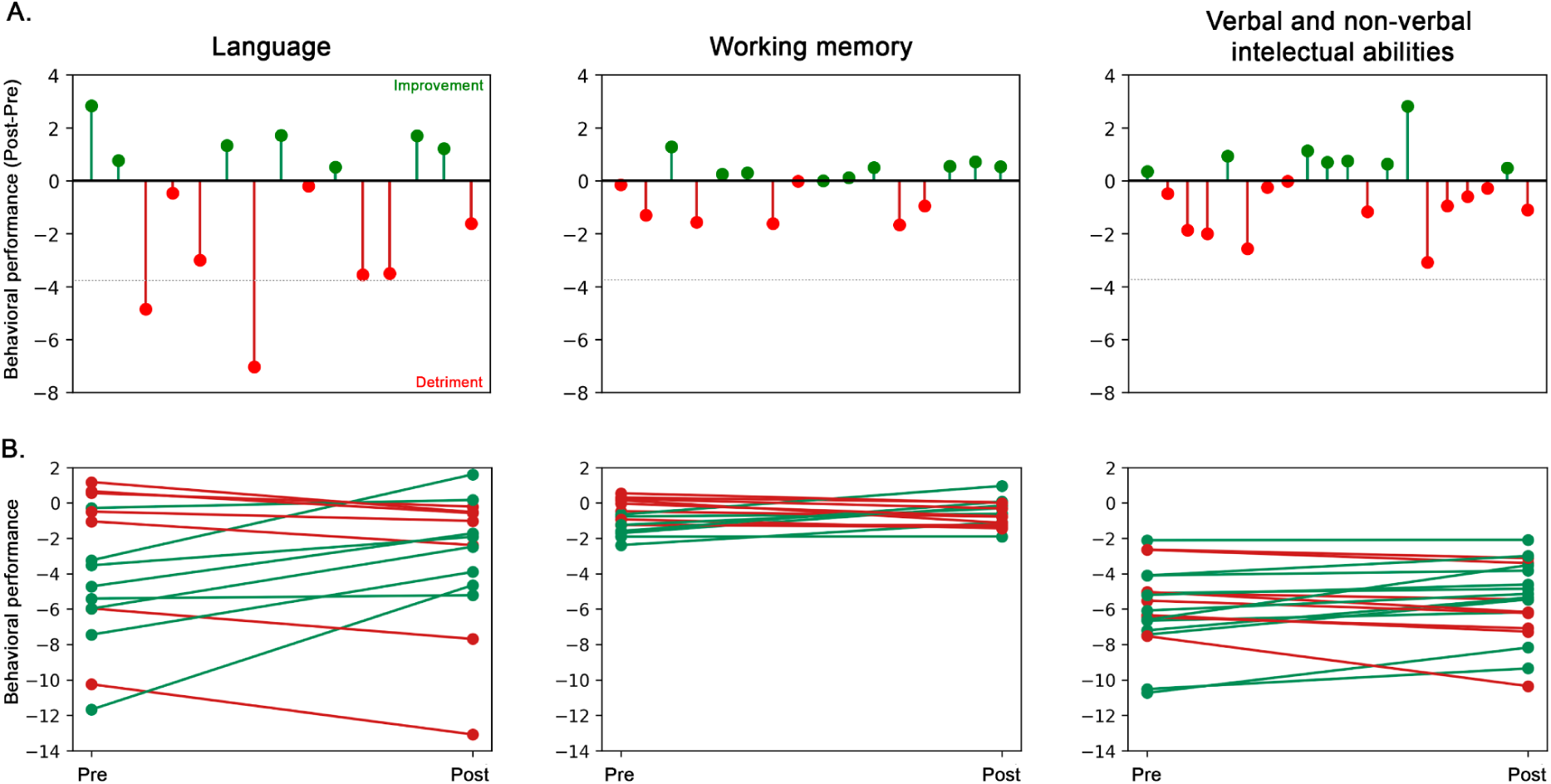
Behavioral results. A) Differences in cognitive performance for each cognitive domain between pre and post-surgical stages. The Y-axis represents cognitive performance, and the X-axis represents the participants. Differences are calculated from composite variables created from z-score values. Positive and negative values represent an increase or decrease in cognitive performance, respectively. Dotes below the grey line denote significant deviations from the performance of healthy participants. B) Progress in cognitive performance for each cognitive domain between pre- and post-surgical stages. The Y-axis represents cognitive performance, and the X-axis represents the time point. Each cognitive domain is calculated as a composite variable created from z-score values. Each participant is represented by an individual line. Green and red lines represent an increase or decrease in cognitive performance, respectively. The mean of healthy participants is represented by zero in the y-axis.

We evaluated whether cognitive changes were associated with volumetric variation within the right hemisphere. We computed robust percentage-bend correlations between each cognitive-progress variable and right-hemisphere intracranial volume. Given the limited sample size, this analysis should be viewed as exploratory and confirmed in larger cohorts. Overall, we found that improvement in intelligence and language was negatively associated with volume increases in the right hemisphere. No significant associations were found for memory. Complete detailed correlation results can be found in Figure 4.6.

**Figure 4.6.**
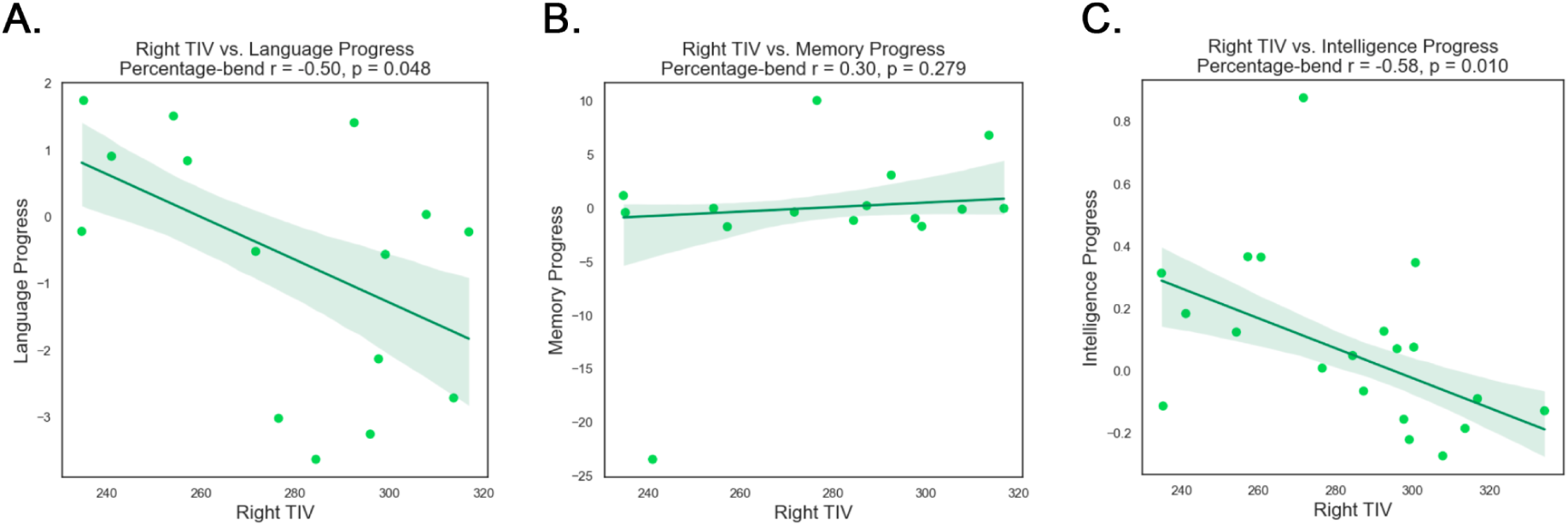
Scatterplots illustrate the relationship between right total intracranial volume (TIV) and key cognitive abilities: language (A), working memory (B), and intelligence (C). Points represent individual data points, and the regression line indicates the linear trend in the data, with shaded confidence intervals.

## Discussion

This study followed patients with left-hemisphere tumors and revealed progressive, widespread structural alterations in the contralateral right hemisphere, offering insights into brain plasticity before and after tumor resection. By focusing on GMV excluding the affected hemisphere and employing both traditional linear regression and data-driven PCA, we identified three key phenomena: 1) significantly larger right-hemisphere GMV in patients compared to healthy participants pre-surgery; 2) an intensification of this divergence post-surgery, evident three months after resection; and 3) a moderate association between these macrostructural changes and post-surgical cognitive performance. These findings suggest that oncological lesions trigger ongoing metaplasticity—higher-order adaptive mechanisms that reshape whole brain architecture far beyond the immediate tumor environment ^6^.

### Macrostructural differences between patients and healthy controls before surgery

Pre-operative enlargement of the contralesional GMV aligns with previous reports of macrostructural alterations in patients with brain tumors ^9,11,19^ and likely reflects early compensatory processes. However, our observation goes beyond replicating these findings by revealing a novel pattern: the non-pathological decline in gray matter volume, widely documented in the aging literature, becomes more pronounced as the onset of the oncological lesion occurs later in life. In the clinical sample, the slope of the age–gray matter volume relationship shifts, showing a steeper decline in patients compared to age-matched controls. This change suggests a complex interplay between adaptive capacity, available neural resources, and aging. Future studies should focus on understanding the role of factors such as prolonged neural stress associated with sustained compensatory processes, considering that the observed relationship may reflect accelerated brain aging induced by the tumor lesion ^36^.

Our PCA based-anomaly detection approach complemented these findings by capturing the multidimensional nature of these structural changes. It revealed that regions that synchronously fluctuate in size are indicative of altered patterns of structural covariance ^33^ and particularly informative in differentiating between the groups. Notably, the olfactory cortex emerged as a primary driver, a region often overlooked in tumor studies but implicated as an early indicator of pathology in neurodegenerative diseases ^37^, potentially due to its role in communicating with higher cortical and limbic structures crucial for cognition ^38^, key to sustaining cognitive abilities. This finding, alongside high reconstruction errors for the hippocampus, amygdala, cerebellum, and temporal poles (superior and middle), underscores a broad response. Indeed, the cerebellum is known for its role in lesion-induced plasticity ^39^, with evidence of cerebro-cerebellar structure-function coupling ^15^ and an increase in GMV reported ^5^. An increase in GMV of the contralesional amygdala has been reported in previous studies in patients with brain tumors ^9^. Further, both the amygdala and hippocampus are critical for interhemispheric communication ^40,41^ and vulnerable to overload from stressors ^42^. Alterations in temporal regions also resonate with the literature on language network reshaping in left-hemisphere tumor patients ^43^, reporting an increase in the superior temporal gyrus ^14^ and decreases in the medial temporal lobe ^11^.

### Longitudinal macrostructural progression

Our longitudinal analysis demonstrated that these GMV alterations are not static, but rather progress at least three months after surgery, leading to further divergence from the normative structural pattern. Key regions implicated pre-surgically, including parts of the cerebellum, superior temporal pole, and amygdala, remained critical discriminators. Additionally, the medial orbital and superior orbital frontal cortices gained prominence post-operatively. The sustained involvement of these regions, known to support higher-order cognitive functions in brain tumor patients in longitudinal ^20^ and cross-sectional studies ^44^, points to a dynamic and enduring neuroplastic response extending well into the recovery phase.

### Relationship between macrostructural changes and cognitive performance

Following the pattern described in previous studies of patients with brain oncological lesions, our patients showed a heterogeneous cognitive profile in the preoperative assessment^45^. Language abilities, as well as verbal and non-verbal intellectual capacities, exhibited notable interindividual variability compared to the healthy control group. The results ranged from patients without detectable behavioral alterations to those with significant impairment relative to the controls’ performance. In the case of working memory, in contrast to what was observed in the other tasks, none of the patients showed significant differences compared to the controls. This finding may suggest a relative functional independence of this domain from the affected left hemisphere.

Postoperative behavioral progression also mirrored the pattern reported in previous studies, with some patients maintaining their performance, while others showed improvement or deterioration compared to the control group ^45^. This differential profile across cognitive domains persisted, with working memory remaining stable between the pre- and postoperative phases. It is particularly noteworthy that, despite these patients having tumors located in the left hemisphere—dominant for language and involving key regions of the network supporting this function—language performance was preserved or even improved in 7 of the 15 cases evaluated longitudinally. Only two patients showed significant deterioration compared to the control group.

Since the PCA based-anomaly detection approach does not allow inference about the directionality of the changes associated with brain damage, an exploratory correlation analysis was conducted between cognitive progression and volumetric changes in the right hemisphere in an attempt to clarify the direction of effects. This analysis showed that these contralateral changes were associated with performance in language and intelligence tasks. Exploratorily, patients with lower increases in contralateral gray matter volume tended to show better cognitive profiles. Although this relationship should be interpreted with caution and does not allow strong conclusions due to the limited sample size included in the longitudinal analyses, these findings, together with the effects detected through PCA, point to a working hypothesis for future studies: research with larger samples should further examine contralateral gray matter volume as a potential predictor of postoperative cognitive decline.

The broader implications of this structure-cognition relationship, particularly whether these GMV changes represent effective compensation, remain an open and critical question in neuro-oncology. Interpreting such associations is challenging. While some studies posit volumetric changes as compensatory, definitive evidence linking specific structural modifications to consistently preserved or improved cognitive performance is scarce. Indeed, the existing literature presents a mixed landscape: for instance, decreased cortical thickness has been linked to agrammatic comprehension ^4^, while increased contralateral frontal GMV in pediatric cohorts was associated with poorer post-surgical cognitive speed and reasoning ^20^, and other work found no clear structural correlates for language abilities ^15^. The common observation of relatively preserved cognition in many patients with brain tumors ^46^ and the potential biases in cognitive assessment practices in the clinic ^47,48^ could be hindering the establishment of clear relationships between changes in structure and behavior.

### General discussion

Our findings support the concept of metaplasticity ^6^ as a pertinent framework for understanding neural adaptations in response to brain tumors. The demonstration of altered patterns of synchronized change in areas far away from the lesion underscores the brain’s interconnectedness and its capacity for large-scale global reshaping. This aligns with previous studies reporting altered patterns of structural covariance in various neurological disorders ^33^, emphasizing the potential of structural co-variance as a biomarker of network-level changes. Data-driven methods, like PCA, can focus on the correlating structure among brain regions rather than specific anatomical measures and can reveal pathologies at the network level. Interpreting these structural covariance patterns, however, remains challenging. Reduced correlations may reflect disconnection between regions, or region-specific changes that are not shared across the network. In contrast, increased correlations could indicate hyperconnectivity or synchronized grey matter loss or gain across distant brain regions.

The debate over whether these patterns reflect actual connectivity is ongoing, as they might relate to the integration or segregation of brain networks in pathological conditions ^49^. This idea needs to be explored through resting-state functional connectivity in the future. Further, the physiological properties of these patterns at the cellular level and their biological implications remain uncertain. Some hypotheses suggest that synaptic changes over time can generate inter-regional structural correlations, which could be tested with animal models ^33^. Finally, patterns of global structural alterations raise the question of whether such changes occur as a consequence or secondary effect of brain tumors as localized lesions, or whether brain tumors—particularly gliomas— should be considered a systemic brain disease^50^. Our results reinforce the notion of brain tumors as a systemic brain disease, emphasizing the brain’s global response to focal pathology.

## Limitations and future research

This study faced several limitations that should be addressed in future research. First, our analysis focused on GMV and did not incorporate surface-based metrics (e.g., cortical thickness, gyrification) or microstructural measures (e.g., from diffusion imaging), which could provide a more complete picture of the structural changes ^51^. Second, although our sample size and inherent patient heterogeneity reflect clinical reality, they limited the statistical power for subgroup analyses and the inclusion of additional demographic or clinical factors. Larger cohorts, potentially achieved through multi-center collaborations, are essential to robustly model these complex relationships and would enable tha application of advanced statistical approaches like Bayesian methods. Third, our cohort was restricted to patients with left-hemisphere tumors who underwent awake brain surgery; investigating patients with right-hemisphere tumors and those undergoing different surgical approaches is crucial to prove the generalizability and potential laterality or surgery-specific effects on contralateral reshaping. The three-month post-operative follow-up may not capture the full temporal evolution of these plastic changes, as structural alterations can progress more slowly than functional ones ^18^. Finally, while we observed different structural covariance patterns, future studies incorporating functional connectivity metrics are needed to elucidate how these large-scale structural synchronies relate to functional dynamics. Addressing these aspects in larger, multi-modal longitudinal studies will be pivotal for a deeper understanding of metaplasticity in brain tumor patients.

## Conclusion

In conclusion, our investigation reveals a widespread pattern of increased GMV throughout the contralesional hemisphere in adult patients with left-hemispheric tumors, suggesting a global metaplastic response characterized by altered structural covariance. These changes, particularly pronounced in regions like the olfactory cortex, amygdala, hippocampus, cerebellum, and temporal poles, not only persist but become more distinct from healthy participants three months post-surgery. Notably, we observed promising moderate correlations between postoperative GMV and cognitive performance, hinting at a potential link between these structural adaptations and cognitive preservation, although these findings require validation in larger cohorts. This study contributes to the understanding of structural neuroplasticity in oncological contexts, highlighting that brain tumor effects extend beyond localized areas.

## Ethics

Data collection was approved by the Ethics Board of the Euskadi Committee and the Ethics and Scientific Committee of the Basque Center on Cognition, Brain, and Language, BCBL (protocol code PI2020022, date of approval: 26 May 2020 and protocol code: 270220SM, respectively). The study protocol was conducted in accordance with the Declaration of Helsinki for experiments involving humans.

## Data Availability Statements

The data are not publicly available due to the data-sharing policies of the different institutions involved concerning vulnerable clinical information. Codes for data preprocessing and analyses will be made available on GitHub upon publication: https://github.com/lmansoo

## Acknowledgments

The authors would like to thank all the participants who agreed to take part in this study and the lab team for assisting with data collection, especially David Carcedo and Maite Kaltzakorta. We are also grateful to Iago Rego García for his valuable insights during our discussions.

## Author contributions

Conceptualization, L.M.-O., I.Q., M.C.; Methodology, L.M.-O., S.G.-M., I.Q.; Software, L.M.-O., S.G.-M., I.Q.; Validation, I.Q.; Formal analysis, L.M.-O., I.Q.; Investigation, L.M.-O., I.Q.; Resources, L.A., G.B., S.G.-R., I.P., M.C., I.Q., Writing—original draft preparation, I.M.-O., I.Q.; Writing—review and editing, L.M.-O., S.G.-M., L.A., G.B., S.G.-R., I.P., M.C., I.Q.; Visualization, L.M.-O., I.Q.; Supervision, I.Q., M.C.; Project administration, L.A., I.Q.; Funding acquisition, L.A., M.C., I.Q. All authors have read and agreed to the published version of the manuscript.

## Informed Consent Statement

Informed consent was obtained from all subjects involved in this study.

## Supplementary Material

**Table 1.**
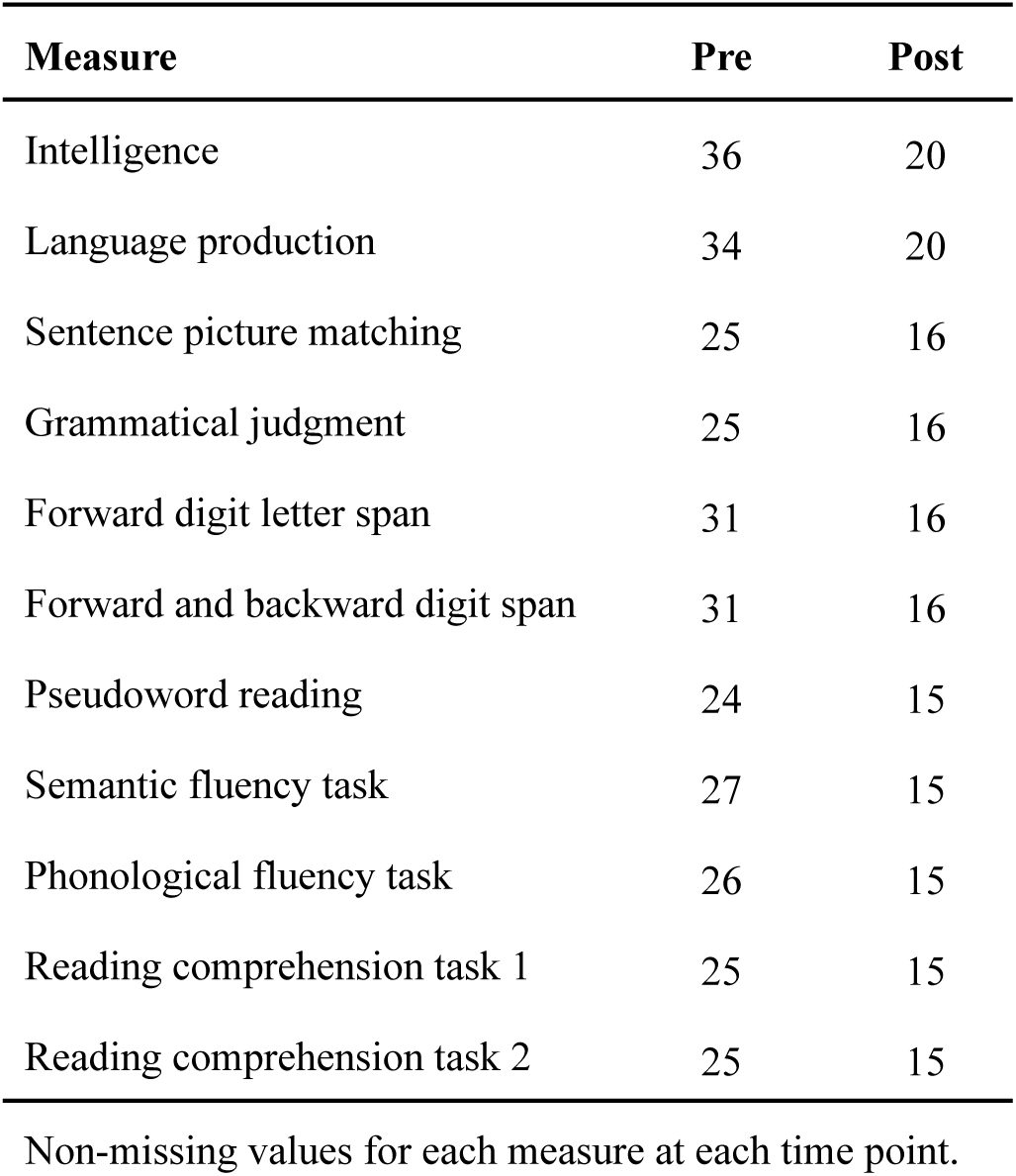
Summary of raw measures per time-point.

**Table 2.**
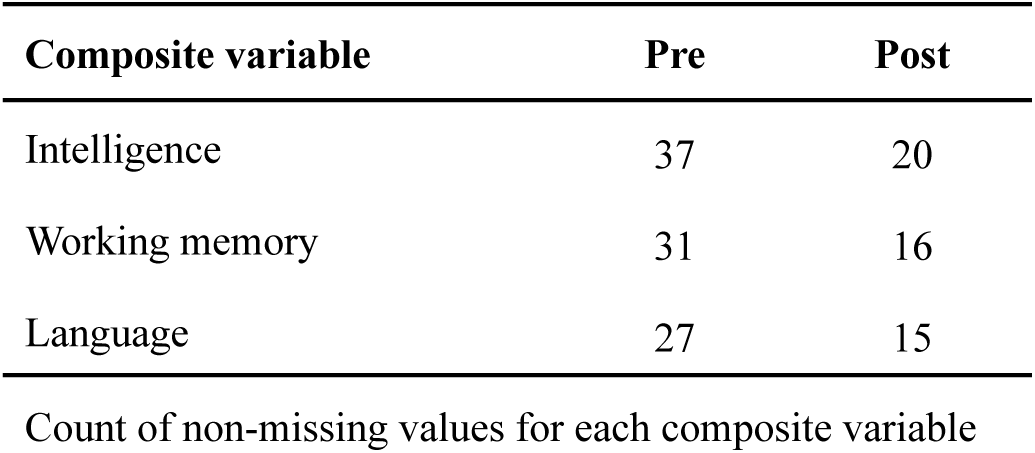
Summary of composite variables.

| Composite variable | Pre | Post |
| --- | --- | --- |
| Intelligence | 37 | 20 |
| Working memory | 31 | 16 |
| Language | 27 | 15 |

Count of non-missing values for each composite variable

