## Supplementary Material for "Contralesional grey matter volume as an index of macrostructural plasticity in patients with brain tumors"

**Table 1.** Summary of raw measures per time-point

| **Measure** | **Pre** | **Post** |
| --- | --- | --- |
| Inteligence | 36 | 20 |
| Language production | 34 | 20 |
| Sentence picture matching | 25 | 16 |
| Grammatical judgment | 25 | 16 |
| Forward digit letter span | 31 | 16 |
| Forward and backward digit span | 31 | 16 |
| Pseudoword reading | 24 | 15 |
| Semantic fluency task | 27 | 15 |
| Phonological fluency task | 26 | 15 |
| Reading comprehension task 1 | 25 | 15 |
| Reading comprehension task 2 | 25 | 15 |
| Non-missing values for each measure at each time point. | | |

**Table 2.** Summary of composite variables

| **Composite variable** | **Pre** | **Post** |
| --- | --- | --- |
| Cognitive functioning | 37 | 20 |
| Working memory | 31 | 16 |
| Language | 27 | 15 |
| Count of non-missing values for each composite variable | | |
